# Nanopore-based consensus sequencing enables accurate multimodal tumor cell-free DNA profiling

**DOI:** 10.1101/2024.02.16.580684

**Authors:** Li-Ting Chen, Myrthe Jager, Dàmi Rebergen, Geertruid J. Brink, Tom van den Ende, Willem Vanderlinden, Pauline Kolbeck, Marc Pagès-Gallego, Ymke van der Pol, Nicolle Besselink, Norbert Moldovan, Nizar Hami, Wigard Kloosterman, Hanneke van Laarhoven, Florent Mouliere, Ronald Zweemer, Jan Lipfert, Sarah Derks, Alessio Marcozzi, Jeroen de Ridder

**Author notes:** To whom correspondence should be addressed: AM; JdR.

## Abstract

Shallow genome-wide cell-free DNA (cfDNA) sequencing holds great promise for non-invasive cancer monitoring by providing reliable copy number alteration (CNA) and fragmentomic profiles. Single nucleotide variations (SNVs) are, however, much harder to identify with low sequencing depth due to sequencing errors. Here we present Nanopore Rolling Circle Amplification (RCA)-enhanced Consensus Sequencing (NanoRCS), which leverages RCA and consensus calling based on genome-wide long-read nanopore sequencing to enable simultaneous multimodal tumor fraction estimation through SNVs, CNAs, and fragmentomics. Efficacy of NanoRCS is tested on 18 cancer patient samples and 3 healthy controls, demonstrating its ability to reliably detect tumor fractions as low as 0.24%. *In vitro* experiments confirm that SNV measurements are essential for detecting tumor fractions below 2%. NanoRCS provides the opportunity for cost-effective and rapid processing, which aligns well with clinical needs, particularly in settings where quick and accurate cancer monitoring is essential for personalized treatment strategies.

## Introduction

A recent and significant advancement in cancer diagnostics involves the high-throughput sequencing of minute concentrations of short cell-free DNA (cfDNA) molecules found in blood and various other bodily fluids (Mandel and Metais 1948). These molecules are primarily released by cells undergoing apoptosis and necrosis but are also actively secreted (Diaz and Bardelli 2014). In patients with cancer, a fraction of the cfDNA molecules stem from the tumor (also known as circulating tumor DNA, ctDNA, or tumor cfDNA). Because these ctDNA molecules carry the genetic features of the tumor (Burnham et al. 2018; McEwen et al. 2020; Husain et al. 2017; Nawroz et al. 1996; Stroun et al. 1987), interrogating them offers exciting opportunities for non- or minimally-invasive cancer screening, cancer diagnosis, minimal residual disease (MRD) detection and monitoring of tumor progression (Lustberg et al. 2018; Bronkhorst et al. 2019; Wan et al. 2017; Corcoran and Chabner 2018; Husain et al. 2022; Diaz and Bardelli 2014). For instance, levels of single or multiple somatic single nucleotide variations (SNVs) can be determined through digital droplet polymerase chain reaction (ddPCR) or targeted sequencing (panels), and several tests based on these principles are currently available (Lustberg et al. 2018; Bronkhorst et al. 2019; Wan et al. 2017; Corcoran and Chabner 2018; Husain et al. 2022; Diaz and Bardelli 2014). However, tumor and MRD detection from small sample volumes (∼10 ml), such as a single vial of blood, remains challenging due to a combination of factors. For instance, the amount of genetic material is in the order of 25ng, corresponding to about 8000 haploid-genome equivalents (Chen et al. 2021; Alborelli et al. 2019). At the same time, the genetic material derived from tumor cells is only a small fraction of this, resulting in extremely low levels of mutated alleles at the targeted positions. Consequently, sequencing artifacts and real mutations may be observed at similar frequencies in raw sequencing data, rendering it very challenging to detect low tumor fractions (Bettegowda et al. 2014; Phallen et al. 2017; Cristiano et al. 2019; Zviran et al. 2020). Finally, subclonal expansions of hematopoietic cells, also known as Clonal hematopoiesis of indeterminate potential (CHIP), is known to contribute to false positive detection of tumor driver mutations and the presence of cancer (Hu et al. 2018; Razavi et al. 2019).

It has been proposed that genome-wide sequencing of cfDNA is a way to circumvent these limitations (Zviran et al. 2020). The likelihood of detecting a tumor is simply enhanced by identifying as many ctDNA molecules originating from a tumor as possible, without the constraints of specific mutations on target or panels of common driver genes. By looking at the mutations through genome-wide ‘breadth’ rather than targeted ‘depth’, the limited number of genome equivalents becomes less restrictive, enabling the identification of tumor fractions at levels approximately 100 times lower compared to targeted SNV analysis (Zviran et al. 2020). Genome-wide cell-free DNA can also reveal tumor-specific copy number alterations (CNAs), which are prevalent in more than 90% of solid tumors and serve as prognostic factors for recurrence (Hoadley et al. 2018; Steele et al. 2022; Martínez-Jiménez et al. 2023; Hieronymus et al. 2018). In addition to traditional genomic mutation features, genome-wide cell-free DNA sequencing enables the detection of cfDNA lengths, cfDNA end motifs, and cfDNA genomic mapping locations, so-called fragmentomics features (Lo et al. 2021; van der Pol and Mouliere 2019; Liu 2021). For instance, cfDNA fragment length typically has a mode of ∼167 base pairs (bp) and a second local maximum at ∼332 bp, related to single or di-nucleosomal DNA (Bronkhorst et al. 2019; Yu et al. 2021). In comparison, cfDNA fragments derived from the plasma of cancer patients show a different length pattern, where fragments are slightly shorter and exhibit a more prominent ∼10 bp periodicity in the fragment length distribution density (Mouliere et al. 2018). Simultaneously evaluating mutational and fragmentomic features in genome-wide cfDNA through a multi-modal approach enables the most efficient detection of ctDNA (Peneder et al. 2021; Moldovan et al. 2023).

A number of sequencing platforms are used for genome-wide cfDNA sequencing, including Illumina short read sequencing, nanopore sequencing and Pacbio sequencing (Lau et al. 2023; Choy et al. 2022; Cristiano et al. 2019), with Illumina sequencing being the predominant platform owing to its low cost per base sequenced. However, both Illumina and Pacbio sequencing typically require substantial initial investments, which may be prohibitive for smaller hospitals and clinical centers in large parts of the world. Moreover, cost-effectiveness is only achieved if samples are processed in large batches, resulting in prolonged turn-around times of several weeks, which is often incompatible with clinical timelines. In contrast, Oxford Nanopore Technologies (ONT) Nanopore sequencing offers low-cost, portable, and non-batched sequencing and is therefore increasingly utilized for cfDNA sequencing (van der Pol et al. 2023; Lau et al. 2023; Marcozzi et al. 2021). Copy number alterations and fragmentomic features can be determined confidently in liquid biopsies with a ctDNA fraction of ∼3% and higher using nanopore sequencing (van der Pol et al. 2023). The main challenge for implementing nanopore sequencing for the purpose of ctDNA sequencing is the relatively high single-nucleotide error rate (Marcozzi et al. 2021). Multiple strategies were proposed to lower the error rate through repetitive sequencing of the same original molecules (Li et al. 2016; Volden et al. 2018; Acevedo et al. 2014; Wilson et al. 2019; Deng et al. 2023; Marcozzi et al. 2021). Whilst SNVs are clearly an important asset in ctDNA analysis, no approach has been successfully implemented to sequence genome-wide SNVs in cfDNA using nanopores (Marcozzi et al. 2021; Martignano et al. 2021; Wilson et al. 2019; Thirunavukarasu et al. 2021; Katsman et al. 2022). Moreover, accurately assessing the presence of a tumor in patients with a low ctDNA fraction remains challenging using nanopore sequencing.

Here we present genome-wide Nanopore Rolling Circle Amplification (RCA)-enhanced Consensus Sequencing (NanoRCS), a high-accuracy genome-wide cfDNA sequencing method based on a combination of rolling circle amplification and consensus calling of long-read nanopore sequences. NanoRCS is a PCR-free approach. Other approaches relying on barcoded PCR amplicons require additional bioinformatics steps to collect copies based on which a consensus is generated (multiple sequence reads yield one consensus sequence), adding complexity and the potential to introduce errors. Conversely, with NanoRCS, cfDNA is amplified using RCA, which minimizes the effects of polymerase errors in the sample since it does not exponentially propagate errors like with PCR. Additionally, the concatemeric RCA products ensure the physical linkage of copies of the same original template, allowing for single-molecule resolution (one native read yields one consensus sequence).

We demonstrate that NanoRCS cfDNA sequencing allows detection of tumor-specific SNVs and CNAs, as well as fragmentomic features. Our proof of concept study spans three cancer types: common, CNA-driven esophageal adenocarcinomas, rare, SNV-driven granulosa cell-tumors (The Cancer Genome Atlas Research Network 2017; Nones et al. 2014; Roze et al. 2020; Smith et al. 2023; Frankell et al. 2019; 2011) and ovarian cancer with different subtypes. We confirm the sequencing-based cfDNA fragmentation patterns using atomic force microscopy (AFM) imaging as an orthogonal and independent measurement (Mouliere et al. 2014). We derived tumor fractions (TF) from all available modalities using computational methods across all cancer samples. We subsequently show that the addition of the SNV modality is especially important in samples with low ctDNA fraction. NanoRCS thus provides an opportunity for cost-effective, sensitive, multimodal cfDNA-based cancer diagnostics.

## Results

### Multimodal genome-wide cfDNA sequencing with NanoRCS

We designed NanoRCS to improve the accuracy of nanopore sequencing, which is especially useful in cases where a low SNV error rate is crucial in the analysis, such as cfDNA sequencing. First, long and short cfDNA molecules are circularized with a specifically designed flexible DNA backbone (Fig. 1A). The optimized circularization reaction and backbone sequences allow cfDNA NanoRCS without PCR and with a cfDNA input of 5 ng obtained from less than 2 ml of whole blood (Alborelli et al. 2019). The circularized DNA molecules serve as a template to create long concatemers by RCA, which are subsequently sequenced on a nanopore device (Fig. 1B). A high-quality consensus sequence of the cfDNA is finally generated from the concatemers using a consensus algorithm to reduce the sequencing errors (Fig. 1C). A true mutation will be present in all repeats, whereas sporadic sequencing artifacts are reduced by the consensus of multiple (≧3) repeats, allowing precise cfDNA sequencing. The entire bioinformatics pipeline is implemented in Nextflow (Di Tommaso et al. 2017) and Snakemake (Mölder et al. 2021) workflow languages, enabling standardized and fast parallel processing of large datasets. NanoRCS offers a multimodal nanopore sequencing-based strategy for cfDNA sequencing through the accurate identification of tumor-guided SNVs along with CNVs and fragmentation length patterns in cfDNA (Fig. 1D).

**Figure 1.**
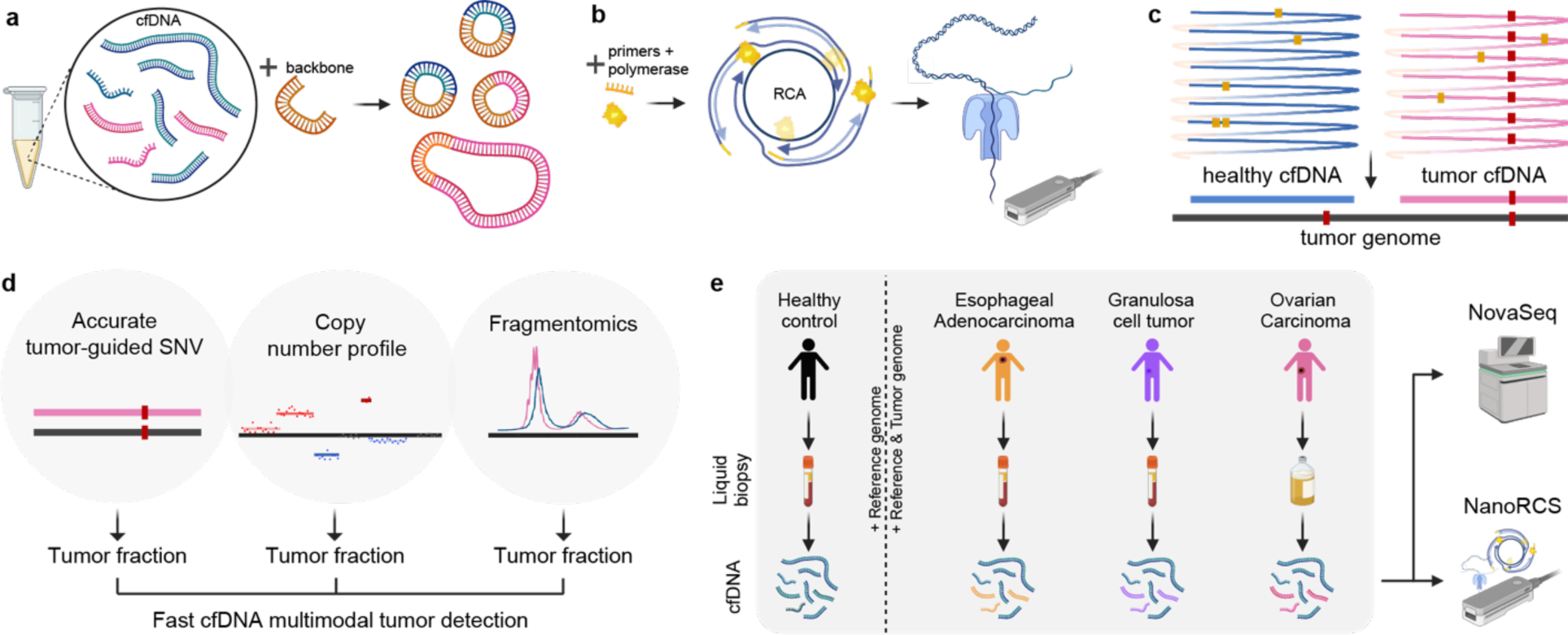
NanoRCS can identify SNVs, CNVs, and fragment size in cell-free DNA. (a) Healthy cfDNA (blue) and tumor ctDNA (pink) are circulated with a high curvature DNA backbone (orange) and form a double-strand circular DNA product. (b) DNA primers of random sequence and phi29 polymerase are added to these DNA circles. Rolling circle amplification of double-strand circular DNA templates subsequently creates long concatemers of cfDNA and backbone that are sequenced on a nanopore device. (c) Obtained DNA sequences are aligned to the reference genome and undergo consensus calling, resulting in cfDNA consensus sequence with reduced random errors (yellow dots) and retained true variants (red dots). After consensus calling, the tumor mutations can be found in the tumor cfDNA and not in the healthy cfDNA. (d) NanoRCS allows simultaneous assessment of fragmentomics, copy number profile, and accurate tumor-guided SNVs. ctDNA fraction can be derived based on all three modalities. (e) cfDNA from plasma of healthy controls, esophageal adenocarcinoma (EAC) and granulosa cell tumor (GCT) patients, and cfDNA from ascites of ovarian carcinoma (OVCA) patients is subjected to genome-wide NanoRCS. We also sequenced the same patient samples with Illumina NovaSeq. Created with BioRender.com.

We performed NanoRCS on cfDNA from plasma of three healthy controls (HCs), five patients with metastatic esophageal adenocarcinoma (EAC), and two patients with recurrent adult-type granulosa cell tumor of the ovary (GCT) (including a time-series of one of the GCT patients), and cfDNA from ascites of seven patients with ovarian cancer (OVCA). Within the OVCA ascites samples, three different subtypes were included, one patient with low-grade serous ovarian cancer (LGS), one patient with carcinosarcoma and five patients with high grade serous ovarian cancer (HGS). (Fig. 1E; Suppl. Table 1). NanoRCS libraries of cfDNA were sequenced on ONT Nanopore MinION and/or the PromethION systems. Finally, to allow comparison with Illumina sequencing, shallow whole genome sequencing with NovaSeq was performed for liquid biopsies of all individuals in parallel.

With NanoRCS, 711,140 unique consensus reads per patient were obtained on average after consensus calling (range 114,105-1,291,239) on the MinION platform. Library complexity calculations of these MinION libraries clearly showed that increased sequencing depth would have increased the amount of data substantially for most samples (Suppl. Fig. 1A-B). On PromethION, 4,998,422 unique reads were obtained on average after consensus calling (range 896,261 - 8,905,143). The analysis of library complexity indicated that a few samples with 5 ng input had started to approach their maximum complexity. However, further sequencing could have yielded additional useful data for the majority of the samples (Suppl. Fig. 1A,C).

### Low SNV error rate with NanoRCS allows SNV-based tumor detection

The main shortcoming of native nanopore-based cfDNA sequencing is the difficulty to accurately detect SNVs. We evaluated if NanoRCS lowered the SNV error rate sufficiently to allow accurate somatic SNV detection in cfDNA (Fig. 2). Germline variants from three HCs were collected, and cfDNA of these HCs were sequenced to determine the SNV error rate of NanoRCS. To calculate the error rate, we selected reads that did not overlap with any single-nucleotide-polymorphisms (SNPs) in the three HCs so that all detected variants could be considered false positives (see Methods section ‘Cell-free DNA SNV detection error rate’). Based on this analysis, we found that the error rate of non-consensus called raw NanoRCS reads was 0.00744 (1 in 134, Q21; Fig. 2A). The consensus calling approach employed in NanoRCS improved the error rate ∼9.3 times lower to 0.00082 (1 in 1,212, Q31; Fig. 2A). This error rate was approximately two times lower than the error rate of Illumina NovaSeq (0.00167; 1 in 598; Q28; Fig. 2A). NanoRCS outperformed NovaSeq even after a consensus sequence was derived from the overlapping paired Illumina reads (0.00108; 1 in 922; Q30; Fig. 2A). Notably, the NanoRCS error profile was very uniform across the genome, i.e. the errors occurred at similar frequencies across all chromosomes, whereas for NovaSeq the GC-rich chromosome 19 (Alan Harris et al. 2020) was clearly enriched for sequencing errors (Suppl. Fig. 2A) (Alan Harris et al. 2020). Performing NanoRCS increased fraction of reads with quality score above Q40 drastically from 31.5% in raw NanoRCS to 88.8% with consensus NanoRCS (Suppl. Fig. 2B).

**Figure 2.**
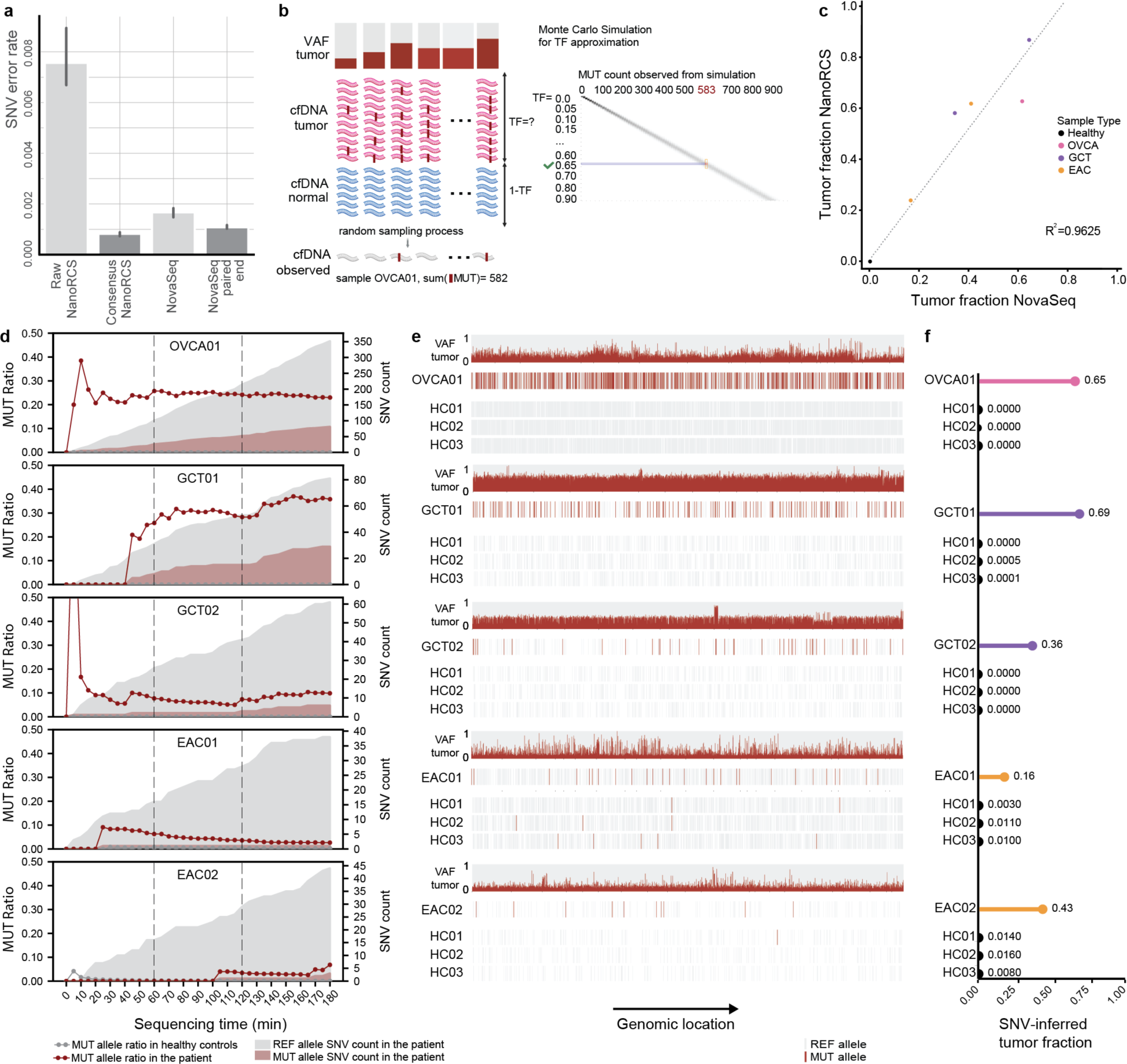
NanoRCS enables SNV detection with a low error rate on Nanopore. (a) Single-nucleotide error rate in cfDNA of three healthy controls using different sequencing methods (the lower the better). Error bars represent standard deviations. (b) Monte Carlo simulations are utilized to search for the tumor fraction that best explains the observations in cfDNA given the known tumor variants and their variant allele frequency (VAF) in the tumor. (For detailed methods, please see *Method section: Tumor fraction estimation from somatic SNV detection*) (c) Correlation of SNV-derived tumor fraction between NanoRCS and NovaSeq. (d) Realtime assessment of the ratio of the mutant (MUT; red shading) and reference (REF; grey shading) allele at somatic SNV positions during the first three hours of sequencing in the five liquid biopsy samples with known somatic SNV profile from the tumor biopsy. Red data points and lines indicate the MUT SNV ratios in cancer patients, and dark grey data points and lines in healthy controls (at level ∼0.00). (e) SNV observations in the five liquid biopsy samples with known tumor somatic SNV profile. For each panel, the top row shows the VAF of detected mutations in the tumor biopsy, the second row represents the MUT or REF allele observations in the liquid biopsy of the corresponding patient, and the bottom three rows represent the observations in three healthy controls randomly downsampled to same amount of observation as in the tumor sample. (f) Lollipop plots show each sample’s inferred tumor fraction corresponding to tumor-informed variants in figure (e) using method described in (b). Created with BioRender.com.

Next, we performed tumor-informed somatic SNV detection in five cfDNA samples where tumor biopsy sequencing is available (Suppl. Table. 2; see Methods section ‘Reference and tumor whole-genome sequencing data collection and preprocessing’ and and ‘EAC reference and FFPE tumor sample whole-genome sequencing’). Tumor-informed somatic SNV detection in cfDNA samples focuses specifically on the sites where mutations have been observed in the tumor biopsy sequencing. The count of mutant alleles versus wildtype alleles were aggregated from the cfDNA molecules overlapping the mutated genomic sites. The mutation fraction was subsequently calculated and compared between each tumor cfDNA sample and three HCs to detect significant elevations above background HC level.

In total, between 10 and 582 somatic SNVs were detected in each of the five tumor samples (Fig. 2C, Suppl. Fig. 2C-G). False positive SNV calls were very low for OVCA01, GCT01, GCT02, at only 0-2 observations per sample, while in EAC samples a higher background of between 5-18 false positive SNV calls was observed (Fig. 2C, Suppl. Fig. 2H-L). In EAC samples, somatic SNVs were determined by sequencing FFPE tumor biopsies which is known to cause false positive somatic SNV calls due to lower sequencing quality and fixation related artifacts (Robbe et al. 2018; Do and Dobrovic 2015). This implies that the majority of the false positive calls in EAC cfDNA may have been, in fact, false positive calls in the tumor biopsies rather than in the cfDNA. Regardless, mutant allele fractions were significantly higher in all patients versus HCs (Fisher’s exact test, p-value <1.2x10^-11^), confirming that NanoRCS can capture tumor-informed somatic SNVs in cfDNA confidently.

### Real-time cfDNA sequencing and SNV-based tumor fraction inferences

ONT Nanopore sequencing uniquely offers the functionality of real-time sequencing of DNA molecules, where real-time results can be analyzed and reported while the during sequencing is ongoing. To demonstrate how this can be leveraged, we also determined the sequencing time required for finding the mutant alleles for each sample. We find that, within 20-110 minutes of sequencing, all 5 samples were above the healthy control background level.

To reliably estimate the TF from SNV data, we developed a Monte Carlo simulation approach. The simulations incorporate the observed number of tumor-informed SNVs, the VAF of each SNV in the tumor and the sequencing error rate. By simulating whether the mutant allele is observed, based on the aforementioned parameters, the most probable tumor fraction is estimated for the tested sample (Fig. 2B; Methods section ‘Tumor fraction estimation from somatic SNV detection’). Based on the Monte Carlo simulations, we determined that the tumor fractions of the patients were between 0.18 and 0.69 (Fig. 2D; Suppl. Table 2). TFs of the patients were more than 36x higher than healthy control background noise (∼0.002; Fig. 2C). The SNV-based tumor fractions observed through NanoRCS and NovaSeq were highly correlated (*R*^2^ =0.96; Fig. 2F), confirming that the SNV error rate is sufficient to estimate the ctDNA fraction in a liquid biopsy reliably.

### CNV-based tumor detection using NanoRCS

CNAs represent another important feature of genome-wide cfDNA sequencing, and can be used to estimate ctDNA fraction. To obtain CNA profiles, we used ichorCNA, which is optimized for CNA analysis on ultra-low-pass genome-wide sequencing data (Adalsteinsson et al. 2017) (Methods section ‘Copy number alteration analysis and tumor fraction inference’). CNAs were observed in 13 out of 14 patient cfDNA samples (Fig. 3A-B; Suppl. Fig. 3A). Six out of seven OVCA ascites samples showed numerous significant increases in copy number, indicative of whole-genome duplications, which often occur in late stage high-grade serous ovarian carcinoma (Cheng et al. 2022; Bielski et al. 2018; Yang et al. 2022). This finding fits with the disease subtypes of these samples, in which five HGS and one carcinosarcoma, and only not observed in the LGS sample, OVCA04 (Fig. 3A; Suppl. Table 2). EAC samples typically exhibited many copy number gains and losses (Fig. 3A), which is in line with the fact that this cancer is typically driven by CNAs (Nones et al. 2014; Frankell et al. 2019; The Cancer Genome Atlas Research Network 2017). SNV-driven GCT (Roze et al. 2020) patients displayed a more copy-number neutral profile, with few CNAs (Fig. 3A). In sample EAC04 we did not observe a clear CNA profile in the cfDNA. This could be due to a low tumor fraction or absence of CNA in this sample. CNA profiles from NanoRCS cfDNA sequencing and NovaSeq sequencing (Suppl. Fig. 3A) highly correlated for most samples (9 out of 14 samples have a Pearson correlation > 0.7, mean = 0.715; Suppl. Fig. 3B), confirming that NanoRCS can capture CNA profiles in cfDNA confidently.

**Figure 3.**
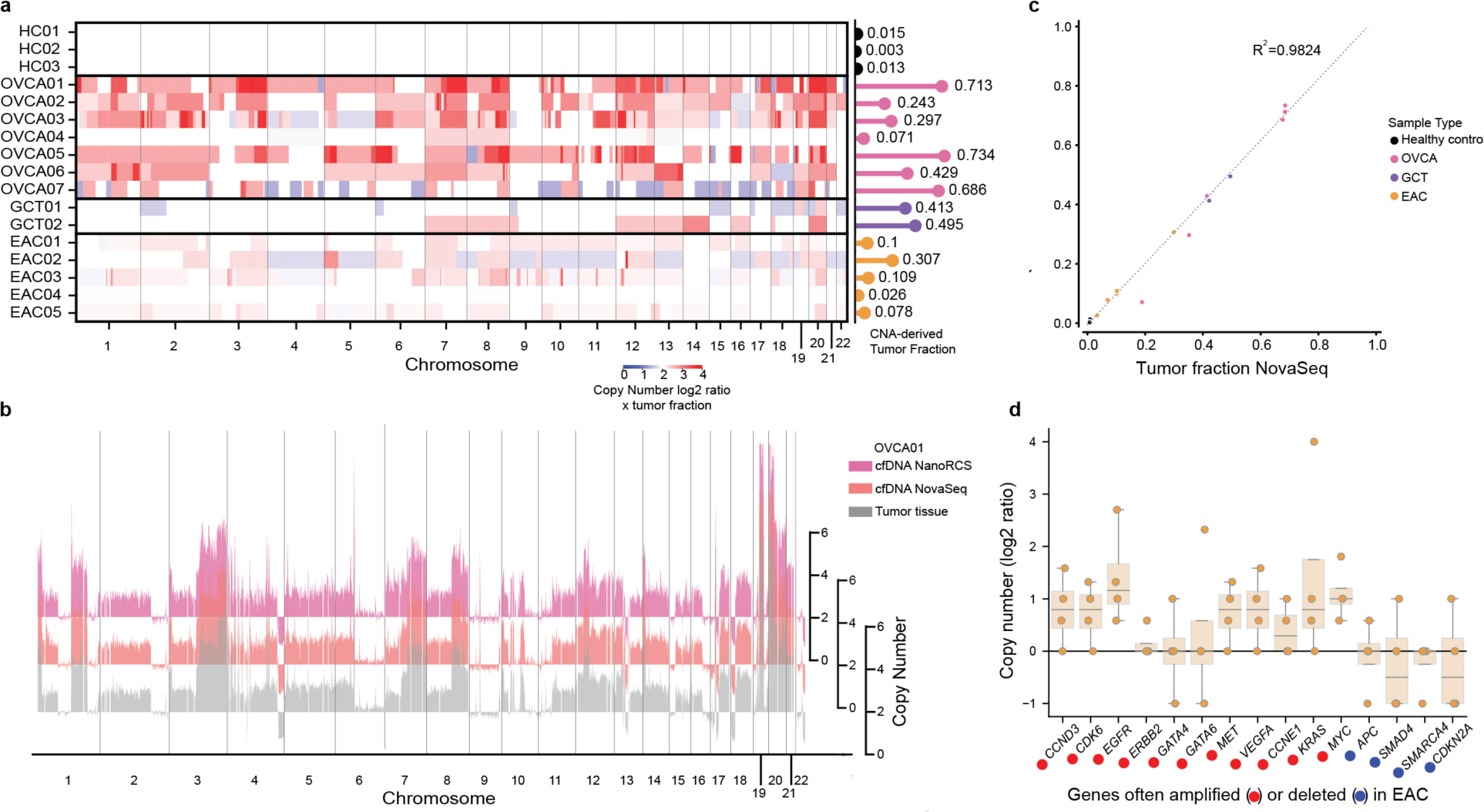
NanoRCS identifies copy number alteration in cell-free DNA of tumor samples. (a) Copy number alterations (CNAs) and derived tumor fraction (lollipops) for cfDNA samples across different sample types (HC, healthy controls; OVCA, ovarian carcinoma; GCT, granulosa cell tumor; EAC, esophageal adenocarcinoma). Red indicates copy number gain and blue indicates copy number loss. Color intensity indicates the copy number alteration multiplied by the tumor fraction in cfDNA. (b) The CNA profile, obtained using NanoRCS on cfDNA (pink), NovaSeq on cfDNA (orange), and from tumor tissue (grey), is concordant in patient OVCA01.The X-axis indicates the genomic position on the chromosomes (1Mb bins), and the Y-axis indicates copy numbers. (c) Correlation of CNA-derived tumor fraction between NanoRCS and NovaSeq. (d) NanoRCS copy number of genes often amplified (red) and deleted (blue) according to literature. The bars show the observed copy number distribution across the four EAC samples in this study. Each data point represents a single (EAC) sample.

The cfDNA CNA profiles were compared to the CNA profiles obtained from the tumor biopsy samples, which were available for five samples. For four out of five samples, CNA patterns were similar in cfDNA and tissue biopsy (Fig. 3B; Suppl. Fig. 3C-G). The more dissimilar sample, GCT02, showed additional copy number changes in cfDNA compared to the sequenced tumor tissue biopsy (Suppl. Fig. 3D). It should be noted that multiple lesions were removed from the abdomen of this patient and only one of them was subjected to WGS. Therefore, the observed difference could be due to differences between the tumor subclones rather than false positive CNA calls (Roze et al. 2020). Indeed, CNA profiles obtained through NanoRCS and NovaSeq were highly similar for this patient as well, suggesting that the CNA profiles were reliable and matched the tumor in general.

ichorCNA infers an estimated tumor fraction simultaneously with CNA using an expectation-maximization algorithm based on a Bayesian statistical framework of a hidden Markov model. In their study, inferred tumor fractions above 0.03 are considered to be reliable (Adalsteinsson et al. 2017). In 13 out of 14 patient samples, we confidently detected ctDNA tumor fractions above 0.03, and tumor fraction of HC samples were below 0.015 (Fig. 3A; Suppl. Table 2). Tumor fractions ranged between 0.07 and 0.73 (median = 0.31). The tumor fraction of EAC04 was too low to be detected reliably with CNA (Fig. 3A; Suppl. Fig. 3A). The OVCA samples, which are ascites samples rather than blood samples, generally displayed high tumor fractions (median: 0.43) compared to plasma samples (median: 0.10). In OVCA01, for which tumor sequencing data was available, the tumor fraction in the cfDNA was even higher than the tumor purity observed in the cell pellet (0.71 in cfDNA and 0.53 in ascites cell pellet). The CNA-based tumor fractions observed through NanoRCS and NovaSeq were highly correlated (*R*^2^ = 0.98; Fig. 3C), confirming that NanoRCS combined with ichorCNA can correctly identify tumor fractions through CNA analysis.

To examine if the CNAs in cfDNA samples corresponded to known driver CNAs in EAC, a list of frequently amplified and deleted genes in EAC was curated (The Cancer Genome Atlas Research Network 2017), and the copy number of these genes in the cfDNA samples was determined (Fig. 3D). From the frequently amplified genes, eight out of eleven are amplified in the EAC cfDNA samples (with the exception of *ERBB2*, *GATA4* and *GATA6*). All patients with EAC in this study have HER2 negative tumors, matching with the neutral *ERBB2* observation (Suppl. Table 2). For frequently deleted genes in EAC, two out of four were found to be deleted in the cfDNA samples as well. We performed similar analyses for CNA driver genes in OVCA and GCT and found that 80.0% and 75.0% of the genes, respectively, that are frequently hit by copy number changes in these solid tumors showed similar patterns in the cfDNA data (Suppl. Fig. 4). The observation of gained and lost genes through cell-free DNA could be useful for detecting driver CNA events, especially in EAC where there are often no SNV-drivers.

### Fragmentomics-based tumor detection using NanoRCS

ctDNA fragments are shorter and exhibit a more prominent ∼10 bp periodicity in the fragment length density compared to healthy cfDNA (Mouliere et al. 2018; Udomruk et al. 2021). In line with this, we found that all plasma samples consistently showed shortening of the first peak, from 169-172 bp in HCs to ∼165-167 bp in GCT and EAC samples (Fig. 4A; Suppl. Table 4). In contrast, samples from OVCA ascites samples displayed variable patterns (Fig. 4A). Some samples did not have an obvious second fragment length peak (sample OVCA02, OVCA04, OVCA06), while others had a high fraction of long reads attributed to this second peak (OVCA03, OVCA05, OVCA07). In addition, there was a more prominent ∼10bp periodic pattern in all OVCA ascites samples. This might reflect the different release mechanisms of cfDNA or differences in cfDNA removal in the abdominal environment compared to the bloodstream.

**Figure 4.**
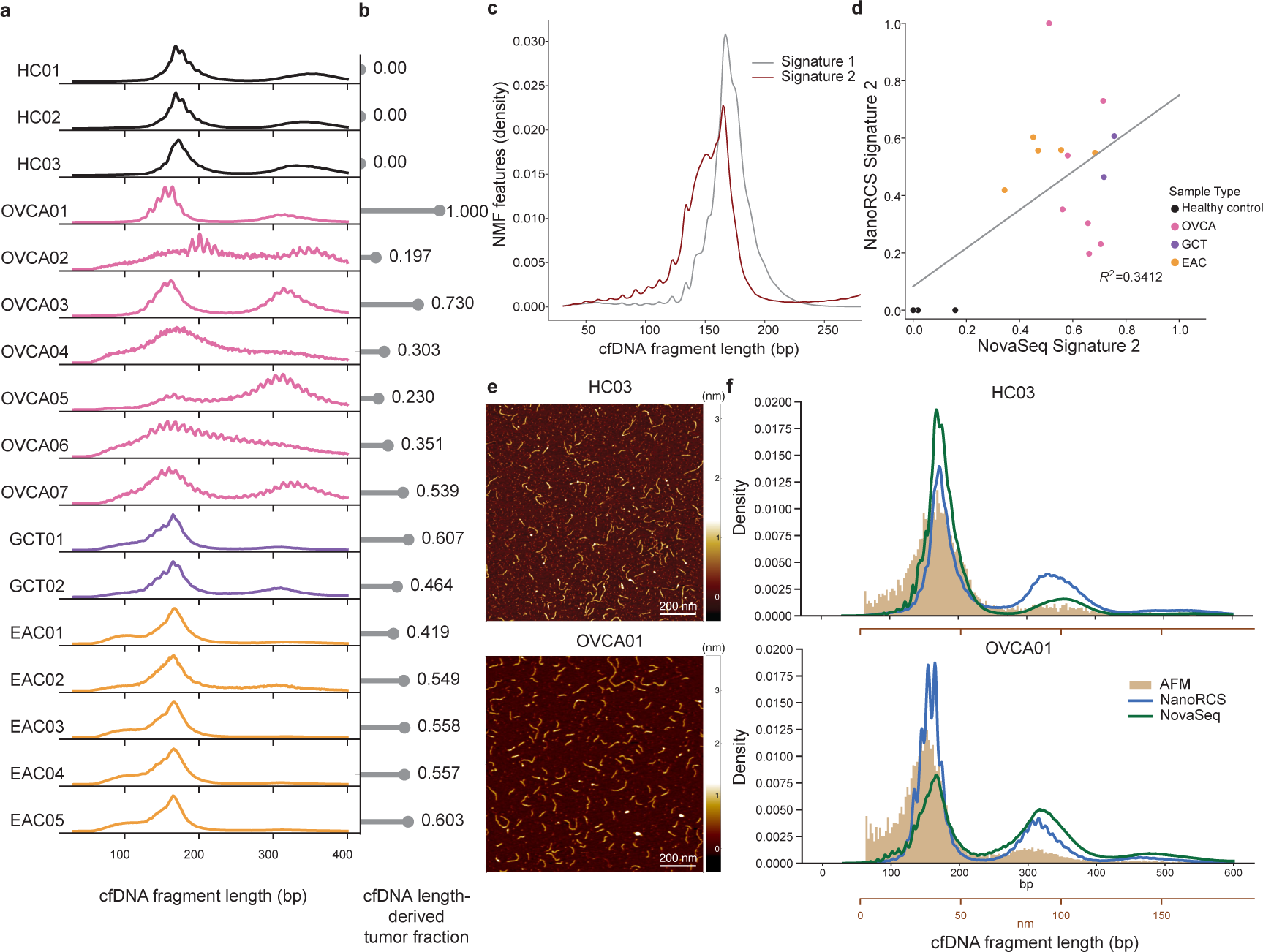
NanoRCS captures cfDNA fragmentation length distribution indicating tumor presence. (a) cfDNA fragmentation length profiles for each sample, categorized by sample type. HC, healthy controls (black); OVCA, ovarian carcinoma (pink); GCT, granulosa cell tumor (purple); EAC, esophageal adenocarcinoma (orange). The profiles display the distribution of cfDNA fragment sizes ranging from 30-400 bp. (b) Tumor fraction (TF) for each sample derived by mapping the length profile to *Signature 2* of the non-negative matrix factorization (NMF) on a reference set (see Methods). TF is annotated to the right of each bar, ranging from 0.14-1.0 for tumor samples. (c) Representative NMF cfDNA length profiles adapted from *Renaud et al.*^55^ for two signatures: Signature 1, predominantly observed in healthy individuals, and Signature 2, indicative of tumor-derived cfDNA. (d) Scatter plot correlating the fragmentomics NMF-derived tumor fraction of NanoRCS and NovaSeq against each other (*R2*=0.3412). (e) Atomic Force Microscopy (AFM) images visualizing HC03 (top) and OVCA01 (bottom) cfDNA fragments (yellow). The scale bar represents 200 nm. (f) Density plots comparing the length distribution obtained from 3 different techniques (blue, NanoRCS; green, NovaSeq; yellow, AFM) in sample HC03 (top) and OVCA01 (bottom). Conversion of the x-axis in nm (brown) to x-axis in bp (black) is based on the calculation of DNA ladder where *L*_bp_ = (*L*_nm_+10) /0.341. AFM imaging (brown) appears to better represent the shorter fragments, while both sequencing methods enriched for longer fragments, especially NanoRCS (blue).

To explore the potential of cfDNA fragmentation patterns as an indicator of tumor presence and tumor fraction in cfDNA samples, we normalized the cfDNA length distribution within the 30-220 bp range and fitted it to a two-component Non-negative Matrix Factorization (NMF) model derived from 86 prostate cancer patients previously sequenced with NovaSeq (Fig. 4B-D) (Renaud et al. 2022; Lee and Seung 1999). Signature 2 in the NMF model recapitulates key features of tumor cfDNA including increased 10 bp periodicity and left-skewed density peak (Fig. 4C). It also closely matches the fragment length distribution of cfDNA with known mutations (Renaud et al. 2022). The contribution of signature 2 therefore served as an indicator of the fraction of cancer-derived cfDNA (Fig. 4B; Suppl. Fig. I-J). The three HC samples did not show any presence of signature 2, whereas the patient samples exhibited contributions ranging from 0.197 to 1.00 (Fig. 4B; Suppl. Table 2). The correlation of NanoRCS-derived and NovaSeq-derived TF was modest (*R^2^*=0.34; Fig. 4D). Taken together, this indicates that, while the fragmentomics-based TF measure may not be very precise, the fragmentatomics-based TF measure does clearly distinguish tumor samples from HCs (all three HCs have derived TF=0).

Sequencing-derived length profiles of cfDNA are affected by DNA purification and bias of the sequencer (Polatoglou et al. 2022; Yu et al. 2023). To verify if our sequencing-based fragment length estimates reflected the true cfDNA length distribution, we employed AFM imaging (Fig. 4E) to measure DNA lengths (Mouliere et al. 2014; Binnig et al. 1986). AFM offers an orthogonal measurement approach based on completely different physical principles, allowing direct visualization of DNA on a flat surface with approximately 1 nm resolution (∼3bp). For selected samples, we took 6-11 images each with an area of 6 μm x 6 μm, for in total between 19,274-107,424 molecules per condition (Fig 4E; Suppl. Fig7 K,M.). Images were analyzed by computationally tracing molecules to obtain AFM-derived DNA length distribution (Fig. 4F; Suppl. Fig. 7). Consistent with the sequencing results, the AFM analysis finds shorter DNA fragment lengths for the tumor samples compared to the healthy control and a relatively larger fraction of molecules in the second peak of the length distributions, especially for OVCA07 (Fig. 4; Suppl. Fig. 7K,L). Comparison of AFM imaging derived cfDNA length profiles to the length profiles obtained by NanoRCS and NovaSeq, suggested that sequencing methods enrich for fragment lengths above 200 bp, while the AFM method was better at detecting shorter fragments (Fig. 4F). NanoRCS was able to capture the second (∼330bp) and third (∼500bp) peak in the fragment length distribution more clearly than NovaSeq. This analysis reveals that while the precise measurements of peak lengths in cfDNA and the proportion of longer to shorter sequences differ based on the method applied, there is a consistent trend of shorter cfDNA fragments in cancer samples as opposed to healthy ones (Fig. 4F).

### Detecting low tumor fractions using multi-modal NanoRCS

By combining multi-modal cfDNA signals, NanoRCS detects disease in all tumor samples and not in healthy samples (Suppl. Fig. 8; Suppl. Table 2). To determine the limit of detection for NanoRCS, we performed variable admixture ratios of tumor cfDNA sample OVCA01 and healthy cfDNA sample HC02 (100%, 10%, 2%, 1%, 0.5%, and 0% OVCA01:HC02 ratio admixtures; corresponding to TF of 0.71, 0.071, 0.014, 0.007, 0.004, and 0.000; Fig. 5A-D). These admixture TFs were derived from estimated TF of 100% OVCA01 sample with CNA modality. TF estimates obtained from fragment length analysis and CNA analysis indicated that TFs down to 0.071 (and above 0.014) could be detected (Fig. 5B-D). The CNA profile at TF=0.071 was consistent with the 100% OVCA01 sample. However, at lower TFs the CNA profile started to deviate from the CNA profile observed in sample OVCA01, suggesting that these CNAs were not reliable. This is in line with the claim of ichorCNA that only TFs>0.03 are confidently detected (Fig. 5B, 5D). Notably, through SNV analysis, the presence of ctDNA was confirmed in all admixtures as low as TF=0.004. The detected TF in the admixture with the lowest ratio (0.004) was 80-fold higher than the background noise observed in HCs. The admixture experiment suggests that genome-wide tumor-informed SNV analysis allows the detection of the lowest TFs, which is required for proper MRD detection (Fig. 5A, 5D).

**Figure 5.**
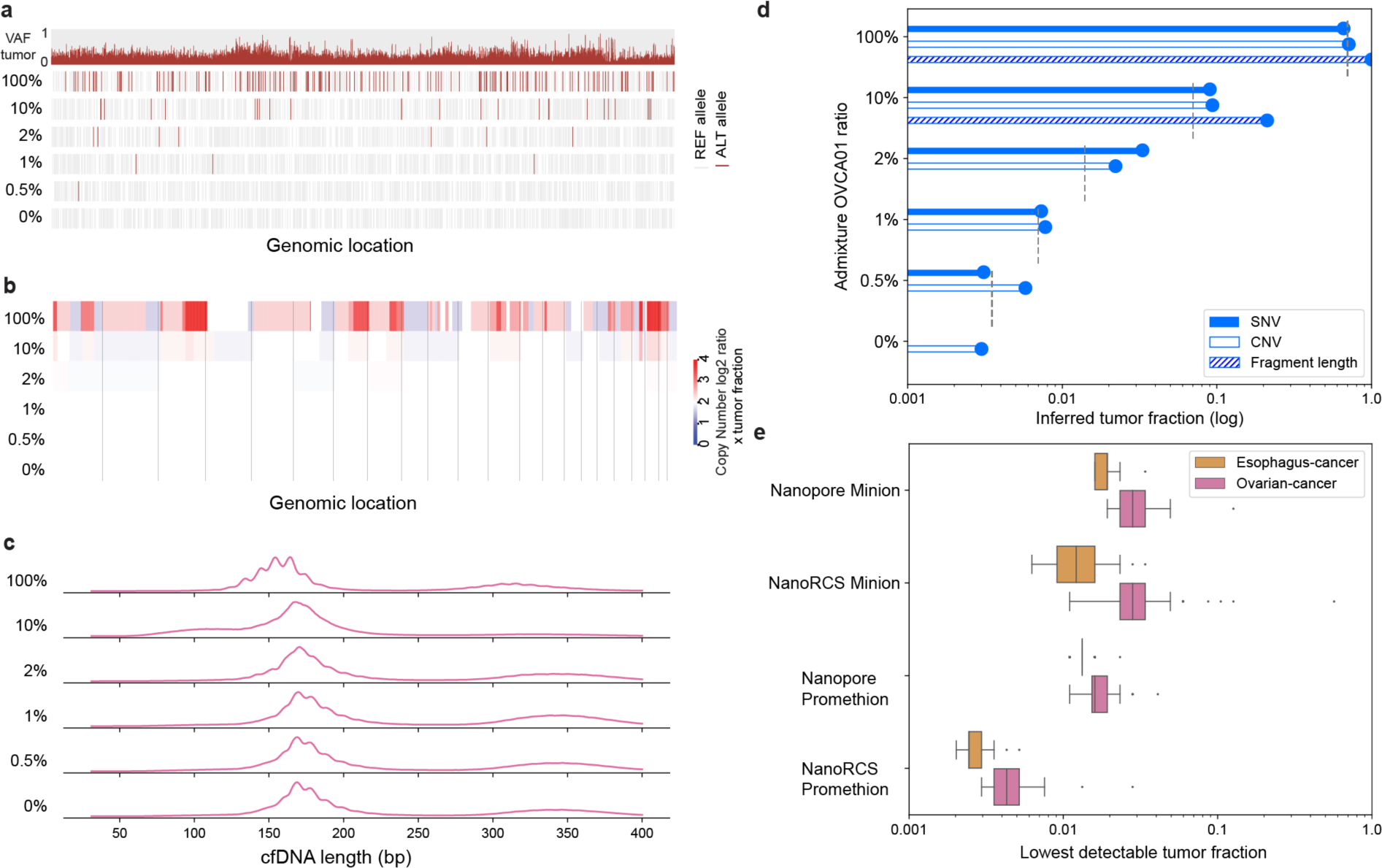
Minimal residual disease detection using NanoRCS. (a-d) Admixture experiment with different percentages of OVCA01 versus HC02 cfDNA (100%, 10%, 2%, 1%, and 0.5%, 0% OVCA01 ratio admixtures). (a) SNV observations for all dilutions. The top row shows the VAF of detected mutations in the tumor biopsy and the following five rows represent the MUT or REF allele observations in the different admixtures. (b) Copy number alterations (CNAs) for all dilutions. Red indicates copy number gain and blue indicates copy number loss. Color intensity indicates the copy number alteration multiplied by the tumor fraction in cfDNA. (c) cfDNA fragmentation length profiles for all dilutions. The profiles display the distribution of cfDNA fragment sizes ranging from 30-700 bp. (d) Inferred tumor fraction of each of the 3 NanoRCS modalities are shown in log-scaled bars. SNV (solid), CNA (nofill), and fragmentation length (striped) are compared to theoretical expected tumor fractions (grey dashed lines). (e) Simulation results to assess the lowest tumor fractions detectable for different platform throughputs (MinION vs PromethION) and with or without consensus calling for Esophagus and Ovarian cancer. Simulation characteristics were obtained from PCAWG (see Methods).

To further deduce the lowest limit of detection, *in silico* Nanopore and NanoRCS MinION and PromethION cfDNA sequencing runs were simulated for 50 EAC and 100 OVCA patients with different variant counts and allele frequencies using the PCAWG dataset at 51 different tumor fractions, each with 10,000 trials (Suppl. Fig. 9A-C) (ICGC/TCGA Pan-Cancer Analysis of Whole Genomes Consortium 2020). These 300 million simulated sequencing runs were generated while taking into account a realistic sequencing throughput (Suppl. Table 1) and error rates (Fig. 2A) of raw and consensus-called MinION and PromethION sequence reads. We then identified the lowest TF detectable for each simulated PCAWG patient in raw versus consensus NanoRCS.

Comparing PromethION to MinION and raw to consensus-called NanoRCS clearly showed that both a lower error rate and a higher throughput could further improve MRD detection (Fig. 5E, Suppl. Fig. 9D). We also observed that higher variant allele frequencies and higher variant counts in samples contributed to improved detection of lower tumor fractions within a cancer type (Suppl. Fig. 9). More than 95% of the samples could be distinguished above background at a tumor fraction of 0.0024 (range 0.0017-0.00535) in esophagus cancer samples and tumor fractions of 0.0044 (range 0.0024-0.0281) in ovarian cancer samples (Fig. 5E).

To further demonstrate the utility of NanoRCS in MRD, we retrospectively applied NanoRCS to cfDNA sampled at five time points of patient GCT02 (Fig. 6; Suppl. Table 2.). GCT02 was diagnosed with adult-type GCT 17 years prior to WGS, and the disease became progressive with unresectable metastases 11 years later. Around the time of cfDNA time series measurements, the patient had multiple chemotherapy treatments and surgery, which led to a temporary stable disease (day 567, 602). The patient, however, relapsed shortly after and died on day 760. For this patient ddPCR-based assessment of the common GCT *FOXL2* mutation was performed at eleven time points (Groeneweg et al. 2021). Over time, disease status and treatment response were collected as well. Fluctuations in ddPCR-based TF estimates were in line with the response on treatments such as surgery or chemotherapy. Using NanoRCS, the TF estimates obtained from all three modalities closely matched the TF values derived from ddPCR at the initial time points. At later time points, however, the TF became very low with SNV measurements both by ddPCR and NanoRCS, while the presence of tumor-derived cfDNA remains evident through CNV and fragmentomics analysis. cfDNA-based CNV analysis revealed emergence of a distinct CNV profile with different gains and losses after the seventh time point (day 497), suggesting that a new subclone becoming predominant. As a result, the known SNV from tumor biopsy was not picked up, while CNV and fragmentomics both showed obvious alteration and increasing inferred tumor fractions (Suppl. Fig. 10). The increased CNA and fragmentomics inferred TFs at late time points corresponded to the patient disease progression at day 720 and death at day 760. This experiment demonstrates that simultaneous evaluation of mutational features (including SNV and CNV) and fragmentomic features in genome-wide cfDNA in NanoRCS enables the most efficient detection of presence of tumor in patients.

**Figure 6.**
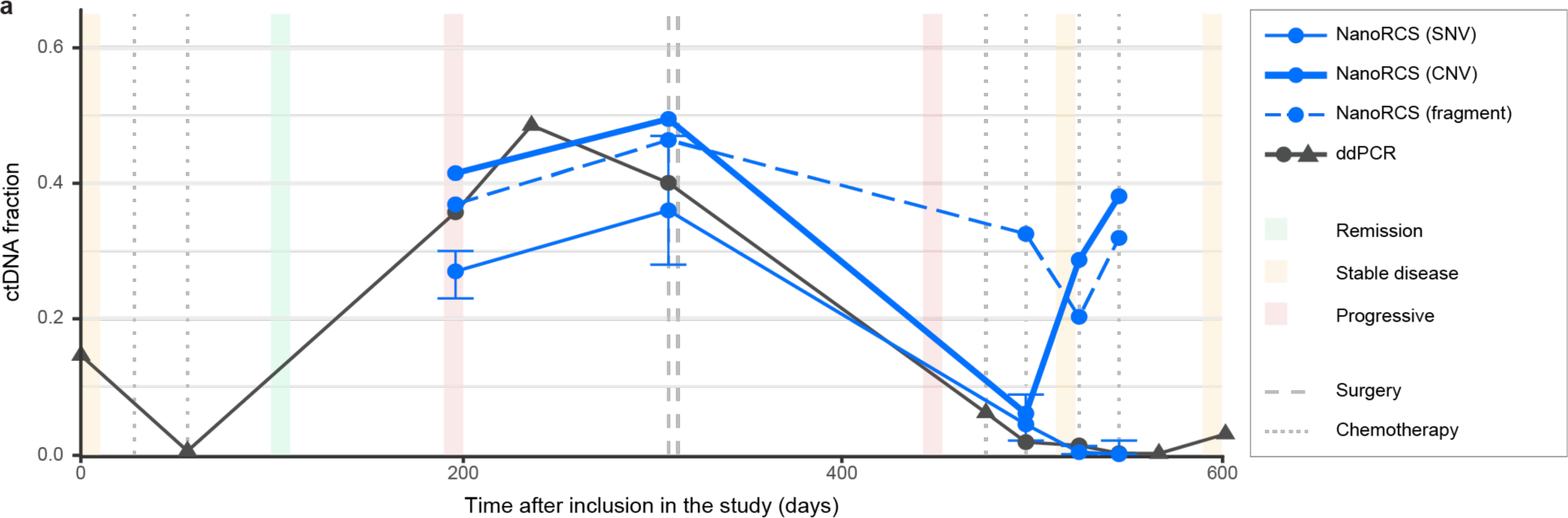
NanoRCS tumor fractions of patient GCT02 during treatment. ctDNA fraction estimations in patient GCT02 obtained using ddPCR (grey) and NanoRCS (blue); the three line types represent the three modalities. A time period of 600 days after inclusion in the study is displayed. Circles represent samples that were analyzed using both ddPCR and NanoRCS, whereas triangles represent samples that were only assessed using ddPCR. Clinical events such as disease state (remission, stable disease, progressive disease) are shown by colored blocks and interventions (surgery, chemotherapy) by vertical lines. Error bars for the SNV modality represent 95% confidence interval.

## Discussion

We introduce NanoRCS, a rapid, highly accurate, Nanopore-based sequencing technology capable of attaining genome-wide cfDNA profiles in a single sequencing run. As shown previously, genome-wide tumor-informed SNV detection in cfDNA can alleviate the limitations associated with detecting one or a few mutation targets in sample volumes with a limited number of genome equivalents present (Zviran et al. 2020). In addition to SNVs, NanoRCS can simultaneously analyze CNAs and fragmentation patterns in cfDNA. As a result, NanoRCS offers a more comprehensive and accurate representation of all subclones present in a tumor, improving the ability to detect tumor presence, monitor tumor progression and identify treatment resistance.

The complementarity of cfDNA modalities is exemplified by our patient time-series analysis and dilution experiments. The TF became (undetectably) low through SNV analyses, both in NanoRCS and ddPCR, while the CNA and fragmentomics modalities provided clear and early evidence for the emergence of a new tumor clone. A distinct CNA profile in late timepoints suggests ongoing tumor evolution, which may go unnoticed when considering single driver events through ddPCR. Tumor evolution is also observed in another metastatic cfDNA sample in this paper with extra CNAs compared to the tumor biopsy. Another example of the complementarity of cfDNA modalities is that we did not observe CNAs nor a CNA-derived TF for one of the EAC samples, while tumor presence was detectable through fragmentomics analysis. Finally, the admixture experiments clearly show that tumor fractions below 3% can only be detected through SNV-analysis. Together, these observations emphasize the significance of capturing multi-modal signals in cfDNA, rather than depending on a single modality.

Lowering sequencing error rate is especially crucial for cfDNA sequencing because real mutations can occur at a similar frequency as sequencing artifacts, making true variants impossible to detect. There is a continued effort by multiple sequencing vendors to lower the single molecule error rate to Q40 and beyond. NanoRCS improved the error rate of Nanopore-based sequencing to the equivalent of Illumina NovaSeq with paired-end error correction, PacBio HiFi reads, and Avidity sequencing (Stoler and Nekrutenko 2021; Arslan et al. 2023; Hon et al. 2020). In addition to NanoRCS, other avenues to lowering sequencing errors are being explored for various platforms. Most notably, amplification with unique read identifiers (UMI) has been demonstrated for both NovaSeq and Nanopore (Karst et al. 2021; Kivioja et al. 2011) as a potent way of producing very high single-molecule accuracies. However, UMI-based methods suffer from clustered amplification errors (Lou et al. 2013). In contrast, NanoRCS is PCR-free and relies on RCA where consensus calls are made from connected copies of the original template, preventing amplification of errors introduced by PCR.

Notwithstanding the importance of lowering the sequencing error rate, our simulations on the efficacy of the SNV modality for tumor presence detection demonstrate that there is a clear trade-off between coverage and error-rate. We conclude that not only low error rate is essential, but a minimal throughput (e.g. 0.8x at Q31) is also required to capture sufficient mutated bases in low tumor fraction samples. This is particularly important for samples and tumor types with low tumor mutational burden and lower tumor stage patients.

An exciting aspect of the ONT Nanopore platform is to deliver real-time sequencing results (Ameur and Hestand 2019; Ardui et al. 2018; Vermeulen et al. 2023; Weilguny et al. 2023). Indeed, NanoRCS can detect ctDNA based on only 20 minutes of sequencing on a PromethION system. Moreover, the ONT platform has experienced continuous improvements in recent years, majorly increased throughput and substantially reduced error rates (Delahaye and Nicolas 2021). These improvements are likely to continue in the coming years, which also directly benefits throughput and error rates of NanoRCS. It should be noted that, although NanoRCS enables more accurate SNV detection in cfDNA, native Nanopore sequencing has the advantage of incorporating methylation as an additional modality that can be captured (Simpson et al. 2017; Lau et al. 2023). Future improvements of NanoRCS could include encompassing even more modalities, such as additional fragmentomic features, to further increase comprehensiveness.

## Methods

### Sample collection

Ascites of patients with ovarian cancer were collected and processed as described in (Kopper et al. 2019; de Witte et al. 2020). The ascites samples used in this study were obtained from the HUB biobank (https://huborganoids.nl/). Plasma of patients with adult-type granulosa cell tumors was obtained previously (Groeneweg et al. 2021). Blood of three healthy donors with available whole genome sequencing data was collected in 10 ml K2EDTA blood collection tubes (BD Vacutainer), and plasma was isolated by two centrifuge steps as described in (Marcozzi et al. 2021). Blood, plasma, and tumors of patients with metastatic esophageal cancer were obtained and processed as described previously (van Velzen et al. 2022). The use of these human specimens for research purposes was approved by the Medical Ethics Committee of the UMC Utrecht (14-472 HUB-OVI, 17-868 and 20/055) and by the Medical Ethics Committee of the Amsterdam UMC (METC 2013_241), respectively. All participants provided written informed consent. Human specimens were stored at -80°C until further use.

### Reference and tumor whole-genome sequencing data collection and preprocessing

Mapped sequencing data (BAM files) of reference blood and tumors from patients were collected from published articles for sample OVCA01, GCT01 and GCT02. For sample OVCA01, data was collected from (Kopper et al. 2019; de Witte et al. 2020). For patients with adult-type granulosa cell tumors, the BAM files were collected from (Roze et al. 2020). Sequencing data of these samples can be found at the European Genome Archive (https://ega-archive.org) under accession numbers EGAS00001003073 and EGAS00001004249, respectively. BAM files of healthy controls were provided by the healthy controls. Since the sequencing data was mapped to different genome versions, BAM files were converted to FASTQ files and then remapped to hg37d5 with bwa mem (0.7.17-r1188) (Li and Durbin 2009), for a uniform data analysis.

### EAC reference and FFPE tumor sample whole-genome sequencing

DNA extraction from metastatic tumor tissue was performed for two EAC patients (EAC01, EAC02) from formalin-fixed paraffin-embedded (FFPE) slides of tumor tissue. DNA was extracted with QIAamp DNA FFPE Tissue Kit (Qiagen) according to manufacturer’s instructions, purified DNA was collected and stored in -80℃ until later use. Matched germline DNA and healthy control from peripheral blood mononuclear cells (PBMC) were collected and stored in -80°C until they were processed for DNA extraction. QIAamp DNA Blood Kits (Qiagen) was used to extract genomic DNA from PBMC. Both FFPE extracted DNA and PBMC DNA Libraries were prepared using KAPA HyperPlus Kit (Roche) following the recommended protocol. Libraries were run on a fragment analyzer to assess their size distribution. Libraries were pooled together based on expected final coverage and sequenced across multiple flow cell lanes. Whole genome sequencing was performed on the Illumina NovaSeq 6000 at 2 x 150 bp read length.

### Somatic variant analysis in tumor biopsies

Whole genome sequencing (WGS) data of tumor biopsies and matched germline samples was mapped to the human reference genome (hs37d5). The WGS data underwent preprocessing steps using the Sarek (Hanssen et al. 2023) nf-core (Ewels et al. 2020) Nextflow pipeline (nf-core/sarek v3.1.2), adhering to the recommended GATK (Van der Auwera and O’Connor 2020) best practices. Default options were activated, including: adapter trimming (Chen 2023) (fastp 0.23.2), alignment (bwa-mem, v0.7.17-r1188), MarkDuplicates (GATK4, v4.3.0.0), and Base quality score recalibration (GATK4, v4.3.0.0). Variant calling was performed subsequently by Strelka (v2.9.10) (Saunders et al. 2012) and Mutect2 (GATK4, v4.3.0.0) with Sarek nf-core Nextflow pipeline by providing paired tumor and normal samples. Single-base somatic variants on autosomal chromosomes with a Strelka filter ‘PASS’ that intersect with Mutect2 results are selected. Subsequently, variants with filter label: “germline” and “panel of normal” were removed to remove potential false positive calls. For EAC samples, an extra step of selecting ‘PASS’ variants in Mutect2 was performed to reduce the amount of false positive calls. With these filtering criteria, a list of tumor-informed somatic SNV were recorded as a VCF file. These VCFs were used in the following Methods section: ‘Tumor-informed SNVs detection in NanoRCS’.

### cfDNA isolation from plasma and ascites

cfDNA was isolated from 5 ml plasma or 5 ml ascites using the Quick-cfDNA Serum & Plasma Kit centrifugation protocol (Zymo Research) for HC, OVCA, GCT samples, and QIAamp Circulating Nucleic Acid Kit (Qiagen) for EAC samples, and then stored in MilliQ at -80°C. DNA concentration was determined by the High Sensitivity Assay Kit (Thermo Fisher Scientific) using a Qubit fluorometer. Between 5 and 25 ng of cell-free DNA was obtained from the human specimens. These cfDNA are used for downstream NanoRCS and illumina NovaSeq sequencing.

### NanoRCS library preparation of cfDNA

NanoRCS is developed in order to generate RCA-derived concatemers of linear and circular cfDNA for subsequent ONT Nanopore sequencing. While circular cfDNA can serve as an RCA substrate as it is, the linear cfDNA needs to be circularized. The circularization reaction used in this study is mediated by a proprietary DNA linker called backbone. Several variations of the NanoRCS were used during this study. Different circularization conditions were evaluated. In particular, we have tested the effect of 1) different amounts of cfDNA input, 2) different backbone sequences, 3) de-phosphorylation of the cfDNA before the circularization reaction, 4) circularization using blunt- and tailed-ends, 5) post-circularization digestion of backbone-backbone concatemers. Each method eventually yielded the same concatemeric DNA as a product, and we did not find any noticeable difference in the consensus reads generated using the different protocols.

The NanoRCS protocol is divided into the following steps:

1. Preparation of the cfDNA for NanoRCS
2. Backbone-mediated circularization
3. Digestion of backbone concatemers
4. Removal of linear DNA
5. Rolling circle amplification
6. De-branching
7. Nanopore library preparation

#### 1) Preparation of the cfDNA for NanoRCS

The cfDNA is prepared to ensure compatibility with the subsequent ligation step. Depending on the protocol variation, the cfDNA can be blunted or tailed, and it can be phosphorylated or de-phosphorylated (Table 1; Suppl. Table 1).

**Table 1.** Protocol variations A-C for preparation of the cfDNA for NanoRCS.

| Protocol | cfDNA ends | cfDNA 5' | Backbone ends | Backbones 5' |
| --- | --- | --- | --- | --- |
| A | blunt | de-phosphorylated | blunt | phosphorylated |
| B | blunt | phosphorylated | blunt | phosphorylated |
| C | G-tailed | phosphorylated | C-tailed | phosphorylated |

For a typical blunting reaction 5-50 ng of cfDNA are mixed with 5 μl of CutSmart 10x (NEB), 5 μl of dNTPs 1mM each, 2.5 μl of DTT and 2 μl of Blunting Enzyme (NEB) in a total reaction volume of 50 μl. The reaction is incubated for 30 minutes at 22°C and heat inactivated at 70°C for 10 minutes. Optionally, the cfDNA can be de-phosphorylated by adding 6 μl of Antarctic Phosphatase buffer 10x (NEB), 2 μl of Antarctic Phosphatase Enzyme (NEB), and 2 μl of water. The reaction is incubated at 37°C for 30 minutes and heat-inactivated at 70°C for 10 minutes. Once blunted, the cfDNA may undergo tailing, using DNA Pol I, Large (Klenow) fragment without exonuclease activity (exo-) (NEB) in combination with dATP or dGTP. The backbone is tailed in a similar fashion to the complementary nucleotide. The cfDNA is then circularized with the help of a backbone during step 2. The reaction is supplemented with three molar excess (relative to the initial amount of cfDNA) of backbone, 18 μl of water, 5 μl of CutSmart buffer 10x (NEB), 5 μl of ATP 10 mM,, and 2 μl of T4 DNA Ligase. The reaction is then incubated at 22°C for 4h and heat-inactivated at 70°C for 30 minutes. When blunt-ended cfDNA and backbone are used during the circularization, the backbone can self-circularize. To remove this byproduct, a Cas9 nuclease digestion is performed. The Cas9 enzyme is loaded with an RNA guide complementary to a sequence formed upon self-circularization of a backbone. In step 3, any residual linear DNA is digested using Exonuclease V (NEB). At this point, only circular BB-cfDNA molecules and original circular-cfDNA molecules are left and will serve as a template for the RCA. During RCA, long concatemers formed by repeats of the circular template are formed. The concatemers are further treated with T7 Endonuclease (NEB) to remove branches before being prepped for Nanopore sequencing.

#### 2) Backbone-mediated circularization

The reaction is supplemented with 3 molar excess (relative to the initial amount of cfDNA) of backbone, 18 μl of water, 5 μl of CutSmart buffer 10x (NEB), 5 μl of ATP 10 mM, and 2 μl of T4 DNA Ligase. The reaction is incubated at 22°C for 4 hours and heat-inactivated at 70°C for 30 minutes.

#### 3) Digestion of backbone concatemers

1 μl of Cas9 nuclease (NEB) and 2 μl of single-guide RNA (check concentration) are added and incubated at 22°C for 30 minutes, then at 37°C for 1 hour. Since Cas9 binding to the DNA can interfere with the next step, the reaction is treated with 2 μl of Proteinase K (NEB). The digestion is carried on at 37°C for 30 minutes, followed by heat inactivation at 98°C for 10 minutes.

#### 4) Removal of linear DNA

The reaction is supplemented with 2 μl Exonuclease V (NEB) and 5 μl of ATP 10 mM, then incubated at 37°C for 30 minutes, then heat-inactivated at 70°C for 30 minutes.

#### 5) Rolling Circle Amplification

Firstly, short random primers are annealed to the template. In particular, 1 μl of exo-resistant RND primers 500 μM (ThermoFisher) are added, the reaction is heated up to 98°C and then let cool down at room temperature. Next, the components for the RCA reaction are added: 22 μl of water, 5 μl of CutSmart 10x (NEB), 2 μl of BSA (NEB), 15 μl of dNTPs 10 mM, 4 μl of Inorganic Pyrophosphatase (NEB) and 2 μl of Phi29 DNA Polymerase (NEB). The reaction is incubated for 8 hours at 30°C.

#### 6) De-branching

The RCA reaction was heat-inactivated at 70°C for 10 minutes, and then 2 μl of T7 Endonuclease (NEB) was added. The reaction was incubated at 37°C for 30 minutes, purified with 75 μl of AMPure XP beads, and eluted with 60ul of H_2_O, pre-warmed at 65°C.

#### 7) Nanopore Library Preparation

Sequencing libraries were prepared according to the manufacturer’s protocol version SQK-LSK109 (for R9.4 flow cells) and SQK-LSK114 (for R10.4 flow cells) using 1500 ng of purified RCA product as input DNA. For barcoding, the Native Barcoding Expansion Kit EXP-NBD109/EXP-NBD114 was used according to the manufacturer’s protocol.

### NanoRCS sequencing and consensus base calling

Prepared sequencing runs are submitted for standard ONT Nanopore sequencing with MinION or PromethION. Sequencing is done via the MinKNOW interface using the SUP base caller model. Multiple fastq files are derived from each ONT Nanopore run. Data is then processed via Cyclomicsseq bioinformatics pipeline automated with NextFlow workflow language to acquire consensus base-calling. In short, ONT Nanopore reads are aligned to the human reference genome (hg37d5) with minimap2. A consensus algorithm takes aligned BAM reads and constructs consensus alignment sequences based on a majority vote for each position, per read. Consensus base quality is calculated by aggregating the quality score for all reads contributing to the final consensus. The remaining backbone sequences are subsequently trimmed from both 3’ and 5’ ends. Reads mapped to the same genomic coordinates are merged.

### Library complexity estimation

An empirical Bayesian methodology by Daley et al.(Daley and Smith 2013) is adopted to determine the molecular complexity of DNA sequencing libraries (shown in Suppl. Figure 1). This technique evaluates the expected unique molecular content in a sequencing experiment. By considering a sequencing experiment as a random sampling from a DNA library, using initial shallow sequencing runs to gauge molecular diversity. Employing principles from capture-recapture statistics, it estimates the frequency of unique molecules, assisting in predicting the yield of additional reads. It uses rational function approximation to enhance accuracy, a technique often applied in theoretical physics. This approach enables precise predictions for sequencing datasets significantly larger than the initial sample.

### Novaseq cfDNA library preparation, sequencing and base calling

Illumina sequencing libraries are prepared by incorporating 1-10 ng of cfDNA with the ThruPLEX® Plasma-seq Kit by Takara Bio, following the guidelines provided by the manufacturer. The quality of these libraries was assessed using the Agilent 4200 TapeStation System, employing the D1000 ScreenTape Analysis assay from Agilent for precise evaluation. Following quality control, the libraries were pooled in equal molar concentrations and sequenced. Two sequencing runs were performed on an Illumina NovaSeq 6000 system, utilizing 150-bp paired-end runs and S4 flow-cells.

Illumina sequencing output is mapped to the human reference genome (hs37d5). Briefly, fastq reads are trimmed with BBDuk (v38.79) before mapping with bwa-mem (v0.7.17). Duplicates in BAM files are removed with Picard MarkDuplicates (picard v2.22.2). BAM files are filtered to remove PCR duplicates, non-primary alignments, and reads with a mapping quality of less than 5. This entire process is automated using a Snakemake workflow language.

### cfDNA SNV detection error rate

SNV detection error rate was established by sequencing cell-free DNA of three HCs with known genome references. Two different sequencing methods were processed in four manners and error rate was compared: raw NanoRCS, consensus NanoRCS, Illumina NovaSeq, and Illumina NovaSeq with paired-end correction. For raw NanoRCS, BAM files are obtained by mapping the ONT Nanopore fastq files obtained with the NanoRCS wet-lab protocol to the human reference genome (hs37d5) with minimap2, settings “-ax map-ont -m 1 -n 10 -s 20” and subsequently keeping the firste alignment of each Nanopore fastq read. Illumina NovaSeq and NanoRCS alignment BAM files preprocessing steps were as described in the sections above. Subsequently, all alignment BAM files are filtered to include only primary alignment reads with a mapping quality of at least 60, then overlapped with the tumor-informed somatic VCF file containing germline variants of three HCs with bedtools intersect (Quinlan and Hall 2010). Reads overlapping with a germline variant in any of the three HCs are removed from the analyses. The remaining reads are supposed to be exactly the same as the reference genome for all samples. The edit distance of each read to the reference is defined as sequencing errors. We randomly subsample 400,000 non-overlapping reads and calculate error rates in all reads. The error rate is calculated by the mismatches excluding indels divided by mapped read length.

### Tumor-informed SNVs detection in NanoRCS

VCF files from germline and somatic variant analysis in tumor biopsies with matched germline samples are acquired as described in Methods section ‘Somatic variant analysis in tumor biopsies.’ These somatic variants specific to the tumors serve as the markers in tumor-informed SNV detection. All mapped reads in the BAM files that overlap with these SNV positions are recorded. The allele overlapping with tumor-informed SNV are noted, reference allele, mutant allele, or erroneous allele (an alternative allele that was not the mutant allele in tumor VCF. Reads overlapping with these identified variants is annotated with timestamp of sequencing for NanoRCS. For each time period, The number and ratio of mutant alleles versus the reference allele are recorded. The number of mutant, reference, and erroneous alleles is recorded for tumor fraction estimation for NanoRCS and illumina NovaSeq.

### Tumor fraction estimation from somatic SNV detection

To estimate tumor fraction from cell-free DNA mixtures, we employed a Monte Carlo simulation approach. Cell-free DNA mixtures are composed of fragments from both healthy cells and tumor cells. Because of the probabilistic nature of observing a mutant (MUT) or reference (REF) allele in tumor-derived cfDNA., each variant position from tumor-derived cfDNA has a probability ***p*** of being observed (adjusted for tumor purity Variant Allele Frequency, VAF); for example, if the tumor purity is 60% and the VAF is 50%, then there is a 30% probability of observing that particular MUT allele. Conversely, there is a ***1-p*** probability of observing a REF allele. For healthy-cell-derived cfDNA, the probability of observing a MUT allele is 0. Therefore, for a given set of MUT alleles and their associated VAFs, we systematically vary the percentage of healthy-cell-derived cfDNA from 0% to 100% in 100 discrete linear steps. For each percentage, we count the number of observed MUT alleles by randomly sampling each allele based on their probability. We repeat this process for 10,000 trials and collect the observed MUT allele frequencies per tumor fraction. The most likely TF in each sample was inferred by identifying where the highest percentage of simulations aligned with the observed distribution of cfDNA sources. The confidence interval is derived from the tumor fractions that fall at the 2.5% and 97.5% of the simulated distribution. If the inferred TF is lower than 5%, we repeated the same process with 100 discrete log steps ranging from 0% to 10% to obtain a more fine grained TF estimation.

### Somatic copy number alteration analysis and tumor fraction inference

We use ichorCNA (Adalsteinsson et al. 2017) software (commit 5bfc03e) to calculate the copy number alterations in all samples. The snakemake pipeline provided is used to perform HMMcopy (version 0.1.1) readCounter to acquire coverage per 1Mb bin and ichorCNA. We modified default settings to allow flexible search for high ploidy (CN=2,3,4) and high copy number (maxCN=8) solutions for all samples and report only chr1-chr22. For samples with inferred tumor fraction below 0.03, we ran extra settings upon the suggestions of the authors on parameter tuning and settings on low tumor fraction samples. Namely, non-tumor fraction parameter restart values were set to lower, and no subclonal status is allowed, with maxCN=3. Detailed parameters provided to the snakemake pipeline can be found in Suppl. table 5. Panel of normal (PoN) files used for NanoRCS runs and tissue biopsy sequencing are constructed with createPanelOfNormals.R. Sequencing of 3 HCs cfDNA and healthy blood cells in this study were used to construct the PoN of NanoRCS and tissue Illumina sequencing respectively. For Illumina NovaSeq runs, PoN provided by authors with ichorCNA software was used. IchorCNA provides several solutions of copy number alterations and derived tumor fraction sorted by the final log-likelihood of the EM convergence. The author stated in the wiki, “*Sometimes ichorCNA will choose a suboptimal solution. Some easy ways to spot a potentially incorrect solution are (1) A large proportion of CNAs are being called subclonal. (2) The majority of data points are brown or red, suggesting a whole genome amplification event. (3) There are two distinct copy number levels being called neutral.”* For low tumor fraction samples (tf = 0.03-0.1), they stated that tumor fraction estimate is most accurate when there is at least one amplification and deletion event spanning more than 100 Mb each. (https://github.com/broadinstitute/ichorCNA/wiki/Interpreting-ichorCNA-results) We used these principles plus the prior knowledge of ploidy of tumor tissue samples to select most likely correct solution for all samples, and the selected solution is provided in Suppl. Tables 6.

### Analysing CNA on tumor-specific copy number events

Tumor specific copy number events are tumor-type specific. Specific copy number events were curated from literature for EAC (Frankell et al. 2019; The Cancer Genome Atlas Research Network 2017), OVCA (de Witte et al. 2022), and GCT (Roze et al. 2020). The list of following genes were curated. For EAC: *KRAS, VEGFA, EGFR, ERBB2, GATA4, GATA6, MYC, CDKN2A, SMAD4, SMARCA4, CCND1, CCND3, CCNE1, CDK6, MET, PTEN, APC, CCNE1*. For OVCA: *KRAS, MYC, CCNE1, TP53, NF1, CDKN2A, RB1, MAP2K4, BRCA1, BRCA2.* For GCT: *FOXL2, TP53, TERT, DICER1*. Genomic coordinates of genes of interest were retrieved from Ensembl biomart version GRCh37. The copy number of the one megabase bin overlapping with these genes was determined to be the copy number of the gene.

### Non-negative matrix factorization derived tumor fraction on fragmentation length distribution

Non-negative matrix factorization is a non-supervised method where a matrix V is factorized into two non-negative matrices W and H, both matrices are smaller than the original matrix V. One of these matrices H, the *signature* matrix, has as many columns as the original matrix and represents the preference of observing each fragment length for each cfDNA source. The other matrix, the weight matrix W, has as many rows as the input matrix and represents the contributions of each cfDNA source to each sample. The number of cfDNA sources is a hyperparameter that needs to be set in advance. Renaud et al (Renaud et al. 2022), utilized NMF for determining the contribution of different cfDNA sources to fragment length signatures in 86 prostate cancer samples. We adapted the 2 signatures extracted from this analysis, and selected the region between 30 to 220 bp, and used them as signatures to decompose the cfDNA source in our sample sets. The fragments between 30-220 bp are selected and normalized to 1. NMF with fixed signatures with function non_negative_factorization from sklearn.decomposition (v1.1.2) is applied to obtain cfDNA source contribution for each sample. All values are capped at 1.0. Implementation for calculating fragment lengths and applying NMF is available (see code availability).

### Atomic Force Microscopy imaging and sample preparation

cfDNA samples and a commercial DNA ladder (Thermo Scientific GeneRuler 50 bp DNA Ladder) were diluted to final DNA concentration of 0.5 ng/mL in buffer of physiological ionic strength at room temperature (10 mM Tris-HCl; pH 7.0; 200 mM NaCl). DNA solutions were deposited by drop-casting a volume of 10 μL (HC03) or 20 μL (DNA ladder, OVCA01, OVCA07) onto poly-L-lysine for 30 seconds, followed by rinsing with 20 mL of milliQ water, and finally dried using a gentle flow of filtered Argon gas (Vanderlinden et al. 2014). The dried sample was immediately measured in air using a Nanowizard Ultraspeed 2 AFM (Bruker), equipped with a high-speed Z-scanner and using a FastscanA cantilever in high-speed tapping mode. A total of 6-11 images of 6 μm x 6 μm were acquired with 4096 x 4096 pixels at a line rate of 3 Hz for each sample.

### Atomic Force Microscopy image analysis for cell-free DNA length determination

Data processing used the software SPIP (Image Metrology, v6.5) and involved background correction using global correction with a 3^rd^ order polynomial and line-wise correction of 3^rd^ degree. The z-offset is set to the mean pixel height after background correction, corresponding to the mean height of the mica surface.

To quantify DNA length distributions from the AFM images, we employed the Particle Analysis pane in Scanning Probe Image Processor (SPIP). A threshold detection level of 0.6 nm with respect to the z-offset was used, and we included a post-processing step to eliminate small speckles with surface-projected area <100 nm^2^. The resulting features were traced by skeletonization and the reported lengths values are the fiber lengths, i.e. the longest segment of a one pixel wide branched line obtained by thinning the 2D-projected surface area of each chain. To test the accuracy and precision of our AFM-based DNA length measurements, we analyzed a DNA ladder sample with fragments of length 50, 100, 150, 200, 250, 300, 400, 500, 600, 700, 800, 900, and 1000bp (Suppl. Fig. 7A-J). Our tracing by skeletonization results in a DNA length distribution for the DNA ladder sample that clearly resolves all 13 fragments in the range from 50 to 1000 bp. The mean lengths determined from the center positions of the Gaussian peaks increase linearly with DNA length in bp (Suppl. Fig. 7C) and the fitted slope of 0.326 nm/bp is in line with previous reports of DNA length from AFM imaging (Konrad et al. 2021a).

Nonetheless, it is noticeable that data points at short lengths fall below the linear fit line, which is readily apparent in a plot of the deviations from linearity (Suppl. Fig. 7D-E), which show small but systematic deviations, with short lengths falling below the fitted constant slope and long lengths above it. These deviations likely stem from the fact that tracing by skeletonization misses some length at the ends of the molecules, which introduces an overall constant offset to the length measurements (since all traced molecules have two ends). We therefore correct the data by adding a constant offset for all values (Suppl. Fig. 7F).

We exploit the fact that the DNA ladder contains DNA fragments in known proportions to test how well our AFM analysis reproduces the expected proportions based on the vendor’s specifications. Overall, the fractions of molecules determined from AFM analyses closely track the expected values, despite the large variation in the number of molecules in each band (Suppl. Fig. 7I-J). The mean (absolute) deviation for DNA lengths < 800 bp is 12%. For long molecules (> 800 bp) the AFM analysis systematically undercounts the number of molecules, which is likely due to the fact that our skeletonization approach excludes strands that cross or self-overlap from further analysis, since the probability to overlap and thus be excluded from further analysis increases with DNA length (Shore et al. 1981; Konrad et al. 2021b).

### Determining lowest TF detectable with generation of cfDNA admixture

We generated cfDNA admixture in the laboratory with samples OVCA01 and HC02. 100%, 10%, 2%, 1%, 0.5%, 0% admixture of OVCA01:HC02 ratio. A 10 times or 100 times diluted stock solution of OVCA01 was used to form a more precise admixture. Final solutions of 5ng DNA cfDNA admixture were subjected to NanoRCS library preparation, followed by sequencing runs with minION and promethION flow cells as described above (and in Suppl. Table 1). Inference of tumor fraction in three modalities including SNV, CNA, and fragmentation length can be found in previous sections.

### Determining lowest TF detectable with simulation on PWACG tumor patient samples

Genome-wide somatic SNV profiles of 50 EAC and 100 OVCA patients were obtained from The Pan-Cancer Analysis of Whole Genomes (Cancer Genome Atlas Research Network et al. 2013; ICGC/TCGA Pan-Cancer Analysis of Whole Genomes Consortium 2020) (PCAWG; https://dcc.icgc.org/pcawg). Real mutation numbers and variant allele frequencies (VAFs) were used as input for the simulations (Suppl. Fig. 9BC). We then generated *in silico* genome-wide sequencing datasets for each patient (Suppl. Fig. 9A) at 51 different TFs (TF of 0 & 50 TFs between: 0.001 - 1, logarithmic) and for 4 techniques: native Nanopore sequencing on Minion (0.25x coverage), native Nanopore sequencing on Promethion (3x coverage), NanoRCS on Minion (0.04x coverage), and NanoRCS on Promethion (0.8x coverage). Error rates for each of the techniques were defined as described in the Methods section ‘Cell-free DNA SNV detection error rate.’ In total, we generated 10,000 simulated dataset for each patient in each scenario, which translates to 2,040,000 (10,000 * 51 (TF) * 4 (technique)) simulated datasets per patient or 3,060,000,000 (2,040,000 * 150) simulated datasets in total. Using the simulated sample with a TF of 0, we could obtain a ‘background noise’ profile for each patient. For each patient, a confident lowest detectable tumor fraction is defined by >95% true positive rate (TPR, i.e. we perform *in silico* sequencing of this patient 10,000 times, and more than 95% of the time we detected the tumor presence) and >68% true negative rate (TNR, i.e. we performed *in silico* sequencing of samples with TF=0 10,000 times, and more than 68% of the time we do not detect cancer presence).

## Data Access

All raw and processed sequencing data generated in this study have been submitted to the European Genome-Phenome Archive (EGA; https://web2.ega-archive.org/) under accession number EGAS00001007475. Raw AFM images were uploaded to Zenodo at doi:10.5281/zenodo.10423114 (HC03), doi:10.5281/zenodo.10423541 (OVCA01), doi:10.5281/zenodo.10423356(OVCA07), doi:10.5281/zenodo.10423726 (DNA ladder) with open access.

## Code Availability

The algorithms and code for conducting analyses in this manuscript is deposited at: https://github.com/UMCUGenetics/NanoRCS/.

## Competing Interest Statement

AM, DM, WK and JdR declare competing financial interests. AM, WK and JdR filed patents and founded a company (Cyclomics) related to the technique. AM, DM, WK and JdR work (partially) at Cyclomics. FM is co-inventor on patents related to cfDNA analysis and has consulted for RocheDX. HvL has consultant or advisory role at Amphera, Anocca, Astellas, AstraZeneca, Beigene, Boehringer, Daiichi-Sankyo, Dragonfly, MSD, Myeloid, Servier Research funding, medication supply, and/or other research support at Auristone, Incyte, Merck, ORCA, Servier and speaker role for: Astellas, Beigene, Benecke, BMS, Daiichi-Sankyo, JAAP, Medtalks, Novartis, Springer, Travel Congress Management B.V.. SD has a consultant or advisory role for BMS (related to checkpoint inhibitors); research funding and medication supply, from Incyte (related to checkpoint inhibitors); and speaker roles for Servier, BMS, and Benecke. The other authors declare no conflicts of interest.

## Acknowledgments

We thank Ivo Renkens of USEQ for the help and advice on ONT Nanopore sequencing, Roy Straver and Carlo Vermeulen for their scientific input on the project, Martin Benoit and the Center for NanoScience at the LMU Munich for help with AFM imaging. We acknowledge the Utrecht Sequencing Facility (USEQ) for providing sequencing services and data. USEQ is subsidized by the University Medical Center Utrecht and The Netherlands X-omics Initiative (NWO project 184.034.019).

## Author Contributions

LC, MJ, WK, AM, JdR conceptually designed the study. LC, MJ, DR, AM, JdR wrote the manuscript. LC, YvdP, NB, AM generated the sequencing libraries. LC, MJ, DR, NM, MPG contributed to data analysis. GB, NH and RZ selected GCT samples and gave clinical input on the GCT and OVCA measurements. TE, HL, SD selected EAC samples and gave clinical input on the EAC measurements. WV, PK, JL performed the AFM microscopy. FM and WK provided expert input on the experiments. JdR coordinated the study. All authors read and approved the final manuscript.

